# Automated cell cycle and cell size measurements for single-cell gene expression studies

**DOI:** 10.1101/182766

**Authors:** Anissa Guillemin, Angelique Richard, Sandrine Gonin-Giraud, Olivier Gandrillon

## Abstract

Recent rise of single-cell studies revealed the importance of understanding the role of cell-to-cell variability, especially at the transcriptomic level. One of the numerous sources of cell-to-cell variation in gene expression is the heterogeneity in cell proliferation state. How cell cycle and cell size influences gene expression variability at single-cell level is not yet clearly understood. To deconvolute such influences, most of the single-cell studies used dedicated methods that could include some bias. Here, we provide a universal and automatic toxic-free label method, compatible with single-cell high-throughput RT-qPCR. This led to an unbiased gene expression analysis and could be also used for improving single-cell tracking and imaging when combined with cell isolation. As an application for this technique, we showed that cell-to-cell variability in chicken erythroid progenitors was negligibly influenced by cell size nor cell cycle.

## Introduction

It has been known for decades that isogenic cells can differ from each other in their molecular composition [1, 2]. The refinement of molecular techniques together with computational approaches has recently allowed to get a quantitative view on this cell-to-cell variability. This strongly highlighted the importance of understanding the causes in such variations, leading to a recent turning point in single-cell studies [3, 4, 5, 6].

A leading source of cell-to-cell variability, or noise, can be attributed to stochastic gene expression, which can be decomposed into “intrinsic” and “extrinsic” sources [7, 8]. The first source can be visualized as the variation in expression levels of identically-controlled genes in a singlecell, whereas the second one affects the expression levels of a given gene between different cells [9]. Intrinsic noise finds its origins in stochastic processes such as diffusion or reactions involving a low-copy number of molecules. It happens especially during transcription and translation processes [7, 10, 11]. Extrinsic noise can be explained by differences in the internal states of a cell population such as cellular age, cell cycle stage or protein subcellular localization [12]. All these factors will contribute to cell-to-cell variability.

It has been shown that stochastic gene expression takes various biological meaning. For bacteria, in a fluctuating environment, generating an heterogeneous cell population can be much more beneficial than an homogeneous population [13, 14]. For HIV virus, the control of stochastic fluctuations determined the latency or the activation of the infection phase [15].

In a cell fate context, stochastic gene expression could drive cells into the differentiation process [16]. For example the lineage choice can be viewed as a stochastic decision process [17]. In line with this view, it has been shown that during the erythroid differentiation process, we can observe an increase in cell-to-cell variability among genes expression that may participate to the decision making process within differentiation [18].

It is thus important to precisely identify the sources of gene expression variability involved in these phenomena in order to understand their role, and to discard potential confounding effects. Flow cytometer FSC parameter can be used as a proxy for cell size, therefore cell size variability in a population can be remove by performing FSC-based cell sorting. Moreover cell cycle-based extrinsic noise can be eliminated experimentally by inducing cell cycle arrest in a specific phase [19] but may result in non physiological alterations. Otherwise cell cycle variability can be identified and suppressed by fluorescent-labeling of cell cycle-specific genes, however this method requires genetical alteration of the investigated cells [20]. Other studies, based on computational approach, deconvolute the cell cycle variables in order to normalize their single-cell gene expression data. Most of them use cell cycle marker genes to train algorithms that can predict cell cycle stage of individual cells [21, 22, 23]. However, these genes have different function or timing according to cell type, even in a same organism [24].

In this article, we propose a more direct approach that consists in measuring morphological parameters in a high-throughtput single-cell RT-qPCR study. Using a non-cytotoxic double-staining technique we measured automatically cell cycle phase and cell size of every single-cell isolated from a primary chicken erythroid progenitor cell population [25]. We demonstrated that the labelling had no detectable effects at the single-cell transcriptomic level in those primary progenitors, suggesting that this technique could be an useful tool for molecular single-cell based studies.

We finally showed that in our cellular system neither cell size nor cell cycle state could be deemed responsible for the cell-to-cell variation we observed, ruling out their potential confounding effects.

## Results

### Cellular morphological automatic measuring

We first attempted to measure cell size manually on C1 images acquired using light microscopy. This proved to be extremely unreproducible and prone to suffer strong experimenter-dependent biases (not shown). To avoid this bias we therefore decided to use cellular as well as nuclear staning.

For this, we choose the two fluorescent dyes, CFSE and Hoechst 33342. CFSE (5-(and 6)- carboxyfluorescein diacetate succinimidyl ester) stably incorporates into cells by coupling both intracellular and cell-surface proteins. It emits high fluorescence intensity and has low toxicity. CFSE concentration is halved with each cellular division, and is widely used to measure proliferation. In this study, it was used as a cell area marker in tandem with Hoechst 33342 [26] as a nuclear marker. The latest rapidly permeates cells and binds to the minor groove of DNA at AT-rich sequences. Like CFSE it can be used without detergent treatment or fixation. Moreover, both markers had been already used for microscopy before performing a RT-qPCR [27, 28]. The use of two different lasers allowed revealing Hoechst and CFSE staining (Figure 1a and 1b) merged in Figure 1c. This double-staining allowed us to automatically measure cell size (CFSE), nucleus size (Hoechst) and the amount of DNA (Hoechst fluorescence intensity). Each individual cell was trapped in the Microchip device. Therefore images were analyzed trap-by-trap in order to eliminate data generated from empty or multi-cell-composed traps. We then automatically retrieved cell size and nucleus size, and inferred their respective volumes.

**Figure 1:**
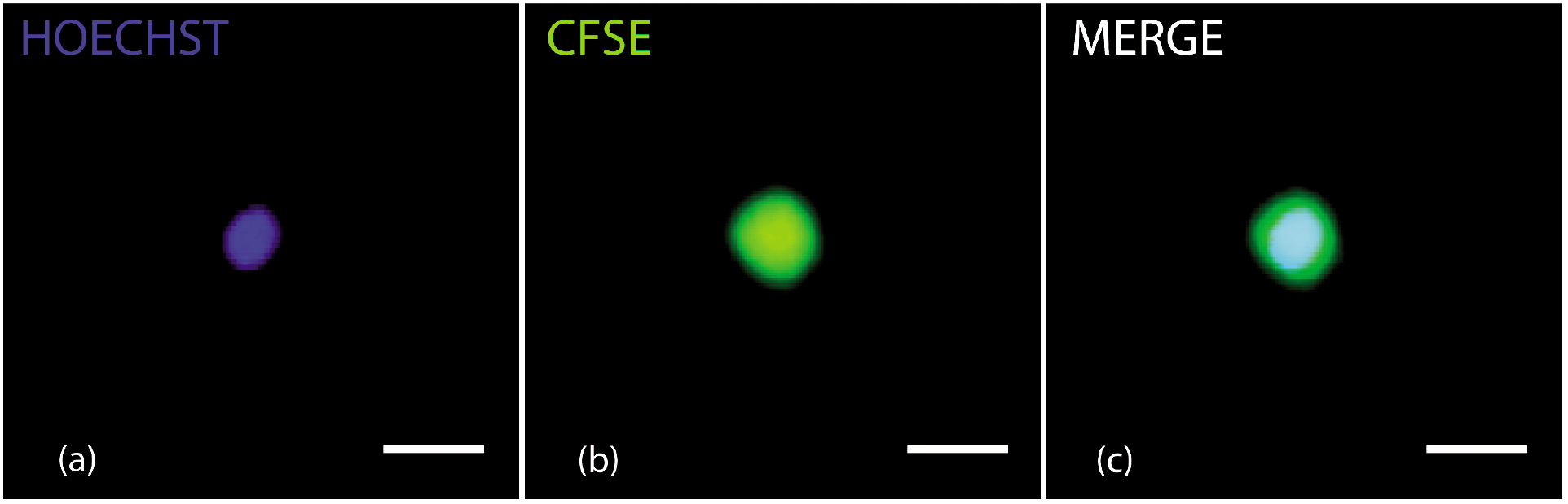
CFSE/Hoechst double staining is compatible with C1 technology. Typical labeling of T2EC nucleus (a) and cytoplasm/membrane (b) stained by Hoechst 33342 and CFSE respectively. (c) Merged image of a and b. Cells were isolated with the C1 system and observed
using a Nikon microscope with 2 different lasers. The scale bar represents 10*μ*m.

We can observe that the cell volume is very poorly correlated with the nucleus volume (Figure 2a). Therefore cell size by itself does not seem to be a good proxy for determining cell cycle position probably because it integrated other unknown parameters. Both cell and nucleus volume density distributions confirm that cell size spans a much larger range than the nucleus size which displays the classical 2n/4n distribution (Figure 2b). Nuclear volume was clearly more correlated with Hoechst fluorescence intensity than cell-volume (Figure 2a & 2c). The nucleus volume can therefore be considered as a good indicator for the position of the cell in the cell cycle. Furthermore it should be noted that volume is a purely geometrical object that does not depends upon intensity and therefore is not influenced by the laser bleaching, as Hoechst fluorescence intensity parameter.

**Figure 2:**
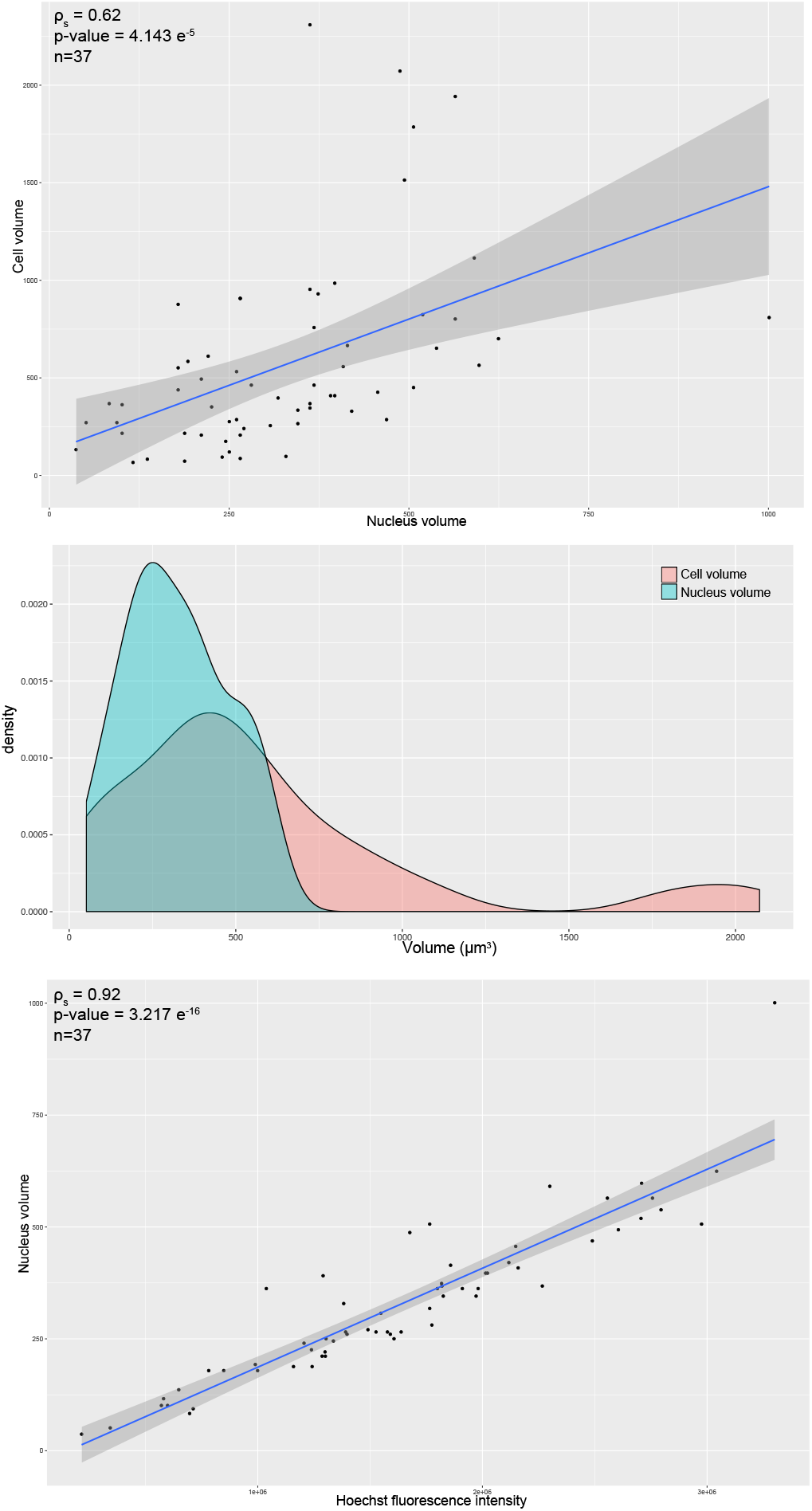
Analysis of cell and nucleus size measurements.

(a) Scatter plot showing the relation between cell volume and nucleus volume. Each point represents a cell. Spearman correlation test was performed, the result of which is displayed in the left upper corner. (b) Distribution of cell volumes (red curve) and nucleus volumes (blue curve). (c) Scatter plot showing the relation between Hoechst fluorescence intensity and nucleus volume. Each point represents a cell. Spearman correlation test was performed, the result of which is displayed in the left upper corner.

We therefore described a double-staining procedure compatible with microscopy associated at the C1 system to measure, for each trapped cell, their size and cell cycle state independently.

### Staining effect

First, we assessed the influence of the double-staining procedure on gene expression at the population level by performing RT-qPCR on 5 selected genes known to be involved in erythroid differentiation or metabolism. The relative value of these gene expressions did not change significantly compared to unstained cells (Figure 3). These results suggested that cell and nucleus staining, with CFSE and Hoechst, had no major influence on T2EC mean gene expression.

**Figure 3:**
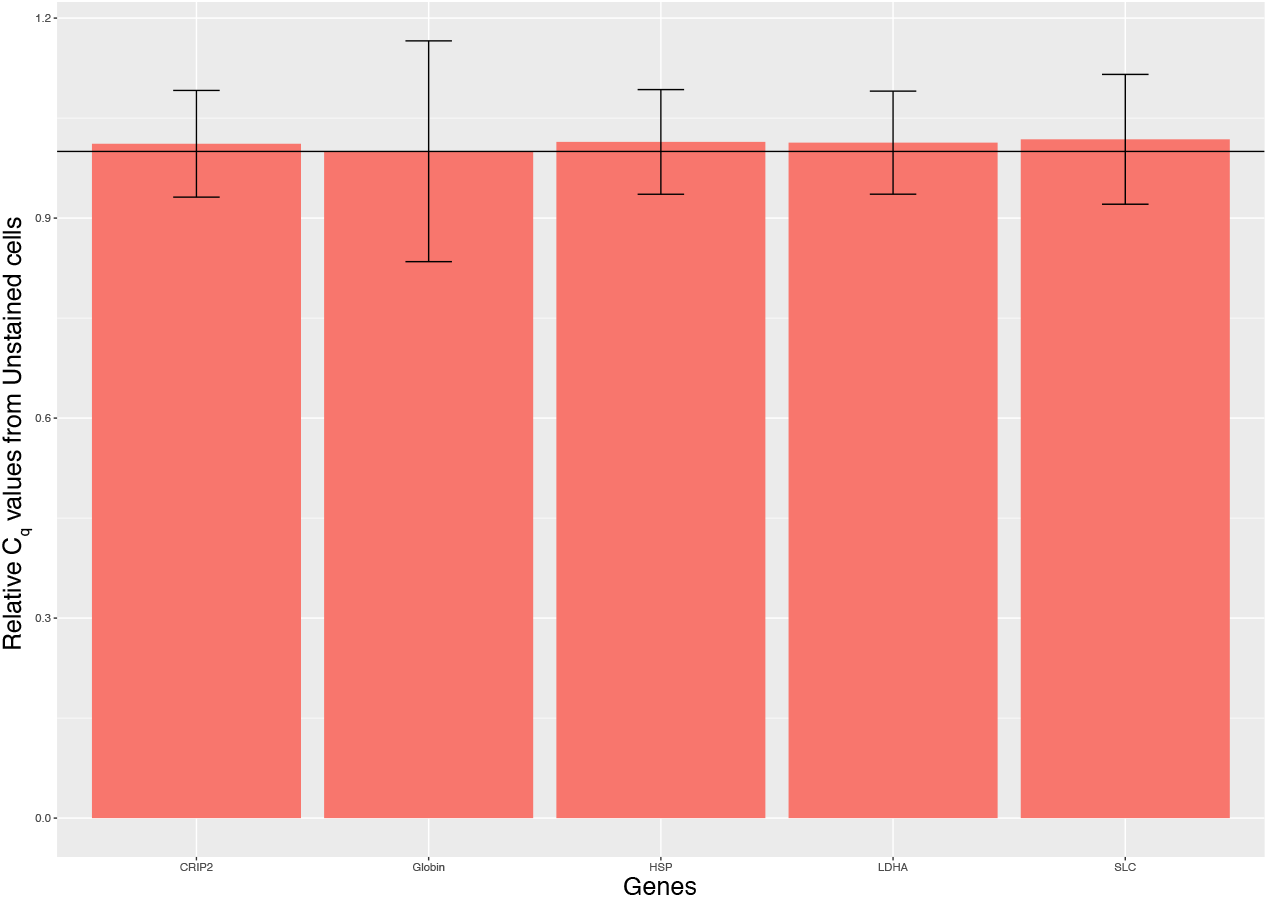
Real-time PCR gene expression analysis of stained and unstained cells. Total RNA was extracted from T2EC cells stained or not. Reverse transcription and real-time PCR analyses, with specific primers [18], were performed to quantify the amount of GLOBIN (*β*-GLOBIN), SLC (SLC25A37), HSP (HSP90AA1), CRIP2 and LDHA mRNA (C_q_ for cycle of quantification). The fold variations represented here correspond to the ratio of mRNA of staining cells compared to unstained cells. The black line corresponds to the null variation between the two conditions. The vertical bars represent the standard error of the mean value (n=3).

In order to extend this double-staining approach to high-throughput single-cell gene expression analysis, we also need to discard possible modifications visible only at the individual-cell level, like cell-to-cell variation in gene expression induced by the staining step.

Therefore we performed high-throughput RT-qPCR on single cells using 92 genes that cover various functions as metabolism, differentiation process and proliferation [18]. We compared 30 single stained cells and 47 single unstained cells in the same microchip. Another plate was done in the same conditions to check the reproducibility. Data was analyzed using a PCA based dimensionality reduction algorithm (Figure 4). The PCA does not show any separation between stained and unstained cells. Moreover, the two first principal components (PC1 and PC2) explained less than 12 % of the variability. These results suggested that the staining did not affect the expression of these 92 genes in T2EC even when examined at the single-cell level. Finally as an application example for our double-staining approach, we investigated the influence of cell cycle and cell size on cell-to-cell variability among our 92 gene expressions using the coupling of labeling and gene expression measurements at the single-cell level.

**Figure 4:**
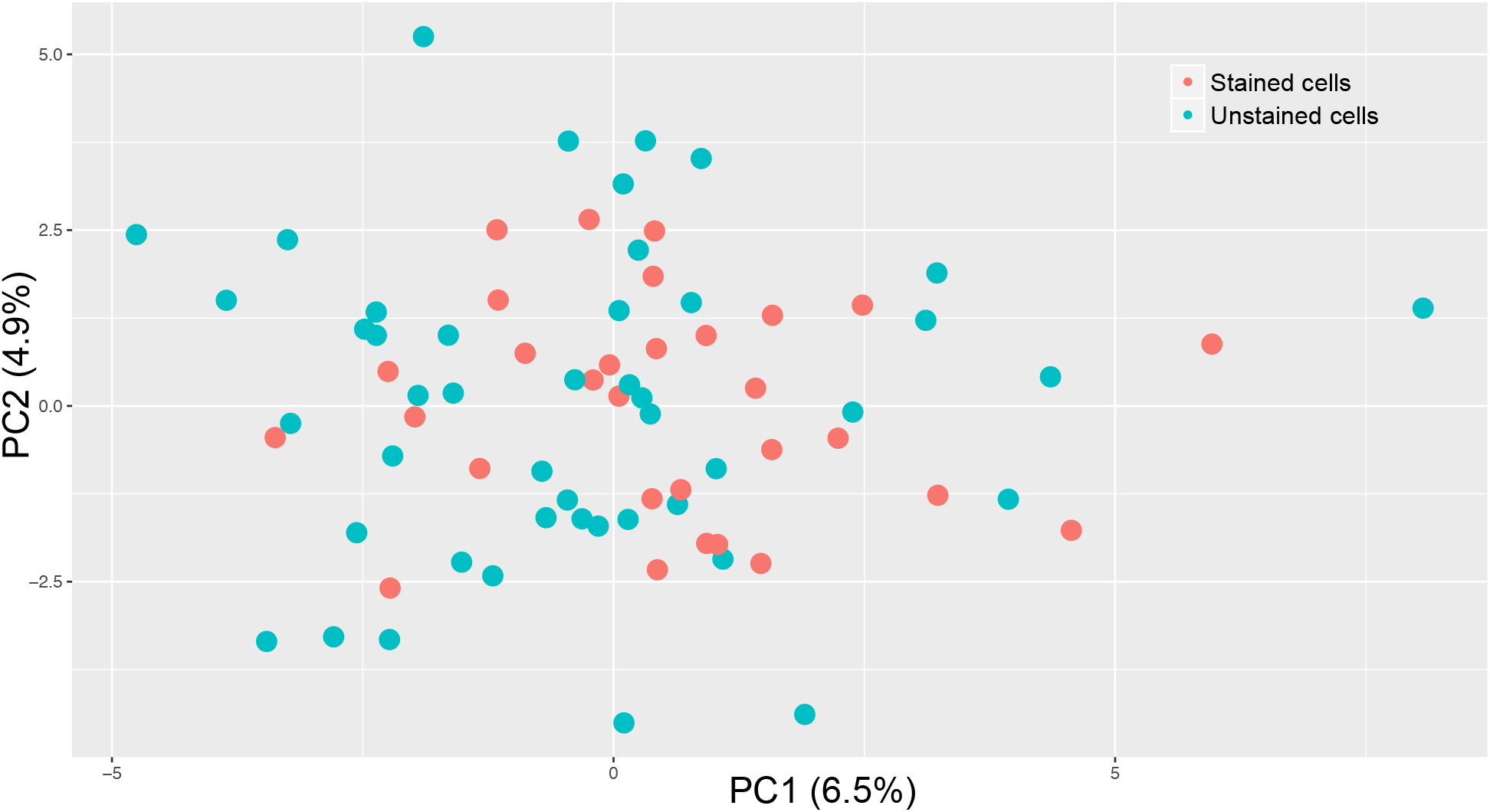
Principal Component Analysis of single cell expression data acquired on stained or unstained cells. Projection of 77 T2EC single-cell stained or not onto PC1 and PC2 results in a cloud of points without any clear separation. Percentages shown are the percentage of variance explained by each component.

### Cell cycle stage and cell size had no impact on T2EC gene expression

For each single cell, we measured the size (CSFE staining), the position in the cell cycle (Hoechst staining) and the mRNA amount (Biomark array). Thereby we could assess whether there were any correlation between morphological and molecular parameters. Among all genes analyzed,none presented a significant spearman correlation between its expression level among single cell volumes or cell cycle : all p-values were above the 5 % threshold. These results confirmed that neither cell size nor the position in cell cycle were relevant parameters in explaining the cell-to-cell variations observed for the 92 genes examined. This information is important for stochastic single-cell-based gene expression analysis, for which these morphological parameters can be excluded of the potential sources of variability between cells.

## Conclusion

In various transcriptome analyses, the question of the influence of cell proliferation on measures have to be assessed [29]. In this study, we propose a way to measure the influence of these factors on a panel of 92 genes without affecting their transcription.

We performed a non-cytotoxic CFSE/Hoechst double-staining compatible with the C1 system. This approach allowed automatic identification and measure of morphological parameters. Used in tandem with the Biomark system, gene expression quantification was then performed. We showed that the double staining did not impact gene expression in our cells. We measured the influence of cell cycle and cell size on a high number of gene expressions. In T2EC, correlation tests between gene expression and these two morphological factors were negative.

The uncorrelated cell size with the total transcript levels has been already shown using flow cytometry sorting of individual cells combined with Biomark system [29].

This method can be used to measure the influence of cell cycle and cell size on single-cell gene expression analysis without any potential misleading cell state effects induced by cell cycle synchronization methods [29, 30]. It could be also an alternative method to avoid artificial cell sorting according to their size or their cell cycle phase, which could be interesting for low amount of cells.

Furthermore, beyond cell size and cell cycle issues, this method can be used to track at least two different cell populations while loading them simultaneously on the same C1 microchip.

## Methods

### Cell culture

T2EC were extracted from bone marrow of 19 days-old SPAFAS white leghorn chickens embryos (INRA, Tours, France). These cells were maintained in *α*-MEM medium supplemented with 10 % Foetal bovine serum (FBS), 1 mM HEPES, 100 nM *β*-mercaptoethanol, 100 U/mL penicillin and streptomycin, 5 ng/mL TGF-α, 1 ng/mL TGF-*β* and 1 mM dexamethasone as previously described [25].

### Double-staining

Cells were incubated in their initial medium for 30 min with CFSE (Life Tech.) at 5*μ*M and Hoechst 33342 (Life Tech.) at 5 *μ*g/mL at 37°C in a tube protected from light. After 2 washings in Phosphate-buffered saline (Life Tech.), cells were loaded in the C1 system (Fluidigm).

### RT-qPCR at population level

Cell culture were centrifuged and washed with 1X phosphate-buffered saline (PBS) 4h after the double staining. Total RNA was extracted and purified using RNeasy Mini Kit (Qiagen).

Reverse transcription assays were performed using the Superscript III First-Strand Synthesis System (Invitrogen) for 500 ng of total RNA.

Real-time PCR was performed with SYBR Green PCR kit (ClonTech) in the CFX96 real-time PCR system (Bio-rad). Specific primers were used to quantify to quantify the expression of genes [18]. For information regarding the specific PCR conditions and primer sequences used, please contact the authors.

### RT-qPCR at single-cell level

#### C1 isolation, capture and RT and pre-amplification

Cells were diluted with C1 cell suspension reagent (Fluidigm) at a concentration of 4.10^5^ cells/mL. This step was followed by a cell filtration in a cellular sieve (50*μ*m). Cells were loaded in the C1 IFC (5-10*μ*m trap size, Fluidigm).

The C1 system performed the cell isolation in each microplate chambers of the IFC. Once cells were isolated, pictures were taken with 2 different lasers (UV laser providing excitation at ∼ 350nm and another laser at ∼ 488nm) using a PALM-STORM NIKON Microscope (CIQLE). This step lasted less than 1 hour before the microplate was back in the C1 system again where lysis, reverse-transcription and pre-amplification was performed. Primers have been previously described [18]. cDNA were loaded in a classic 96 well plate and conserved at −20°C until the RT-qPCR.

#### Biomark PCR

Pre-amplified cDNA were mixed with Sso Fast Evagreen Supermix With Low ROX (Bio-Rad) and DNA binding dye sample loading reagent (Fluidigm). Primers used for pre-amplification were prepared by pair at 5 *μ*M with the Assay Loading Reagent (Fluidigm) and low EDTA buffer. An IFC Controller HX (Fluidigm) performed the prime of a 96.96 DynamicArray IFC Chip (Fluidigm). Then, prepared cDNA and primer pairs were loaded in the inlets of this microfluidic-based chip. Each condition (stained and unstained cells) was loaded in parallel in the same microfluidic-based chip to avoid chip-to-chip technical variability.

The IFC Controller HX performed the load of cDNA samples and primers from the inlets into the chip. The Biomark HD analyzed the chip according to the GE 96 x 96 PCR + Melt v2.pcl program, thanks to the Data Collection software.

RNA spikes were used as positive control to validate the RT-qPCR experiment.

From this outlet, the Real-Time PCR Analysis software generated cycle of quantification values (C_q_) for each reaction chamber (each cell for all primers).

### ImageJ analysis

Each image corresponding at each lasers used were analyzed following a previously described procedure [31]. We visually confirmed the capture for each well and extracted automaticaly morphological information using ImageJ. First, we thresholded the image to highlight cells. Secondly, we applied the analyze particles of ImageJ with correct morphological parameters. ImageJ selected automatically cells. After checking that all cells were detected by the software, we run the measurement of cell area (for CFSE stain), nucleus area and intensity (for Hoechst stain). The cell-volume (2) was then calculated from area measurements (1) using these following formulae:

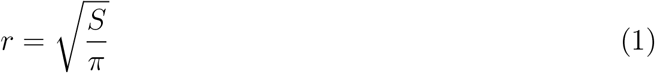

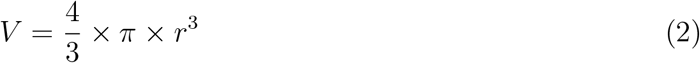

with r the radius of cell, S the area and V the cell volume in *μm*^3^.

### Analysis of gene expression

For population RT-qPCR analysis, ratios of gene expression variation between conditions were calculated following this following formulae [32] :

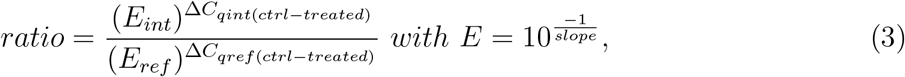

the PCR efficiency of gene of interest E_int_ or referential gene E_ref_.

For each pair of primers, the efficiency of PCR was determined using a standard curve generated by a dilution series of the samples.

ΔC_q_ represents the difference between the C_q_ of control condition (ctrl) and the C_q_ of treated condition for the gene of interest ΔC_q_int or for the standard gene ΔC_q_ref. Because of its low variability between all conditions, HnRNP was used as referential gene in these analyses.

For single-cell RT-qPCR, raw C_q_ data was then computed using R [33] via a specific script that was previously described [18]. The output file comprising absolute values of mRNA was used as a template for all following analysis. Statistical correlations were performed using spearman tests with Bonferonni correction for multiple tests.

### PCA

PCAs were performed using ade4 package [34]. PCA was centered (mean substraction) and normalized (dividing by the standard deviation). PCA was displayed according to PC1 and PC2, which are the first and second axis of the PCA respectively.

## Acknowledgements

This work was supported by funding from the French agency ANR (ICEBERG; ANR-IABI-3096) and La Ligue Contre le Cancer (Comité de Haute Savoie). We thank ProfilXpert platform for the use of C1 system and Denis Ressnikoff (CIQLE) for imagery platform.

